# Maroon rice genomic diversity reflects 350 years of colonial history

**DOI:** 10.1101/2024.05.06.592698

**Authors:** Marieke S. van de Loosdrecht, Nicholaas M. Pinas, Jerry R. Tjoe Awie, Frank F.M. Becker, Harro Maat, Robin van Velzen, Tinde van Andel, M. Eric Schranz

**Author notes:** These authors contributed equally to this work (Tinde van Andel, M. Eric Schranz).

## Abstract

Maroons in Suriname and French Guiana descend from enslaved Africans who escaped the plantations during colonial times. Maroon farmers still cultivate a large rice diversity, their oldest staple crop. Maroon oral history and written records by colonial authorities provide contrasting perspectives on the origins of Maroon rice. Here, we integrated genomic ancestry analyses of 136 newly sequenced Maroon rice varieties with ethnobotanical and archival research to reconstruct the historical contexts associated with the introduction of rice varieties to the Guianas. We found that a large subset traces to West Africa, linked to the transatlantic slave trade (c.1530-1825). Maroons obtained other varieties from indentured laborers from Java (1890 onwards), USA rice breeders (1932 onwards), and Hmong refugees from the Vietnam War (1991). Furthermore, we found rice types never documented before, indicating Maroon farmers selected from crosses. Overall, our results demonstrate that the Maroon farming system prioritizes maintenance of a high stock diversity, which we posit reflects the expertise they inherited from their (African) ancestors. Ignored by agricultural modernization initiatives, Maroon farmers today are important custodians of a unique cultural heritage. Moreover, the genomic findings underline many Maroon stories about their past. This study hence demonstrates the power of cross-disciplinary crop research to reconstruct aspects of the human past for which historical records may be biased or incomplete. We anticipate that a similar study approach can be applied to other heirloom crops of (Indigenous) communities that may have preserved their history on their farms to reconstruct, acknowledge and honor the past.

## Introduction

Marronage, the temporary or permanent escape from slavery, was a widespread act of resistance against the plantation economies in the Americas during colonial times (1500-1870). Today, Maroon communities continue to exist in Suriname and French Guiana, accounting for c. 239,000 people, divided into six distinct groups (Price 2018). Maroon farmers, almost all of them women, have continued to grow various African staple crops using shifting cultivation methods (Fleury 1994; van Andel, van der Velden and Reijers 2016). These crops have remained central not only to their nutrition but also to the preservation of their spiritual and cultural practices (Price 1991; Van Andel, et al. 2019).

A crop of great significance to the Maroons is rice. They grow a large number of rice varieties, sometimes up to twenty-one per farmer (Pinas, et al. 2023). Maroon farming is done primarily by women, who obtain almost all their planting stock via exchange with other Maroon women (Van Andel, et al. 2019; Pinas, et al. 2023). Maroon rice varieties are landraces: crops that are locally adapted by farmers (Zeven 1998). Landraces retain eco-geographical relevant information in their genomes that has allowed for reconstructing crop dispersal and domestication in the past, such as for wheat (Scott, et al. 2019), rice (Gutaker, et al. 2020), and tropical fruits (Zerega, et al. 2004; Larranaga, et al. 2021). Since the Green Revolution in the 1950s, a significant proportion of landraces has disappeared from farmer’s fields worldwide, resulting in severe genetic erosion (Zeven 1998). Many Maroon landraces retain adaptive traits of wild rice that make them more resilient to ecosystem changes and predation compared to commercial cultivars, such as shattering seeds, red pericarp, and long awns (Pinas, et al. 2023). Maroons consider some of their rice landraces as African, and their oral history has preserved numerous stories that provide context to the naming of their rice varieties and origins (Carney 2005; Pinas, et al. 2023; van Andel, et al. 2023).

Here, we leverage the potential of the global rice landrace diversity presented by the 3,000 rice genome (3K-RG) project (The 3,000 rice genomes project 2014; Wang, et al. 2018) to clarify from where various Maroon rice varieties were introduced to the Guianas. The 3K-RG project distinguished nine major genomic groups among the global varieties of Asian rice (*Oryza sativa* L.), including its two subspecies *indica* and *japonica*, most of which are broadly associated with specific (eco-)geographical regions (Han and Xue 2003; Agrama, et al. 2010; Wang, et al. 2018). Ancestry characterization of Maroon rice varieties in this global context of landrace diversity may clarify where in the world their closest genomic relatives are located. By integrating genomic ancestry with ethnobotanical field studies and archival research we aim to reconstruct the historical contexts associated with the introduction of rice varieties to the Guianas and reconstruct events of agronomic innovation in the Maroon past.

## Results

We obtained whole-genome resequencing data for 136 rice varieties collected from Saamaka and Okanisi Maroon communities across Suriname and French Guiana (Figure 1A and S1). In addition, we built a comparative dataset that includes ∼1,400 globally distributed *O. sativa* landraces selected from the 3K-RG project (Wang, et al. 2018), early commercial cultivars from the USA (registered: 1905-1963) (Vaughn, et al. 2021), varieties of African rice (*O. glaberrima* Steud.) (Meyer, et al. 2016; van Andel, van der Velden and Reijers 2016), and relevant diploid wild *Oryza* species (AA genomes) from Africa (*O. barthii* A. Chev., *O. longistaminata* A. Chev & Roehr), and the Americas (*O. glumaepatula* Steud.) (Figure 1A). Subsequently, we called genotypes for the newly sequenced varieties for SNPs overlapping with the 3K-RG 404k coreSNP Dataset (available via https://snp-seek.irri.org/_download.zul) and intersected the Maroon rice data with the comparative data set (Figure S2). After SNP- and individual-level filtering, our final genotype dataset for population genomic analyses consists of 109,203 SNPs (‘∼110k panel’) that are highly ancestry informative for a total of 1,636 rice varieties (Figure S2, Data S1A and B). Our sequenced Maroon rice varieties cover 95.0 ± 5.0% (mean ± s.d) of the SNPs with an average genotype depth of 10.3 ± 6.1X, which overlaps with that for the 3K-RG landraces (14.4 ± 6.9X).

**Figure 1.**
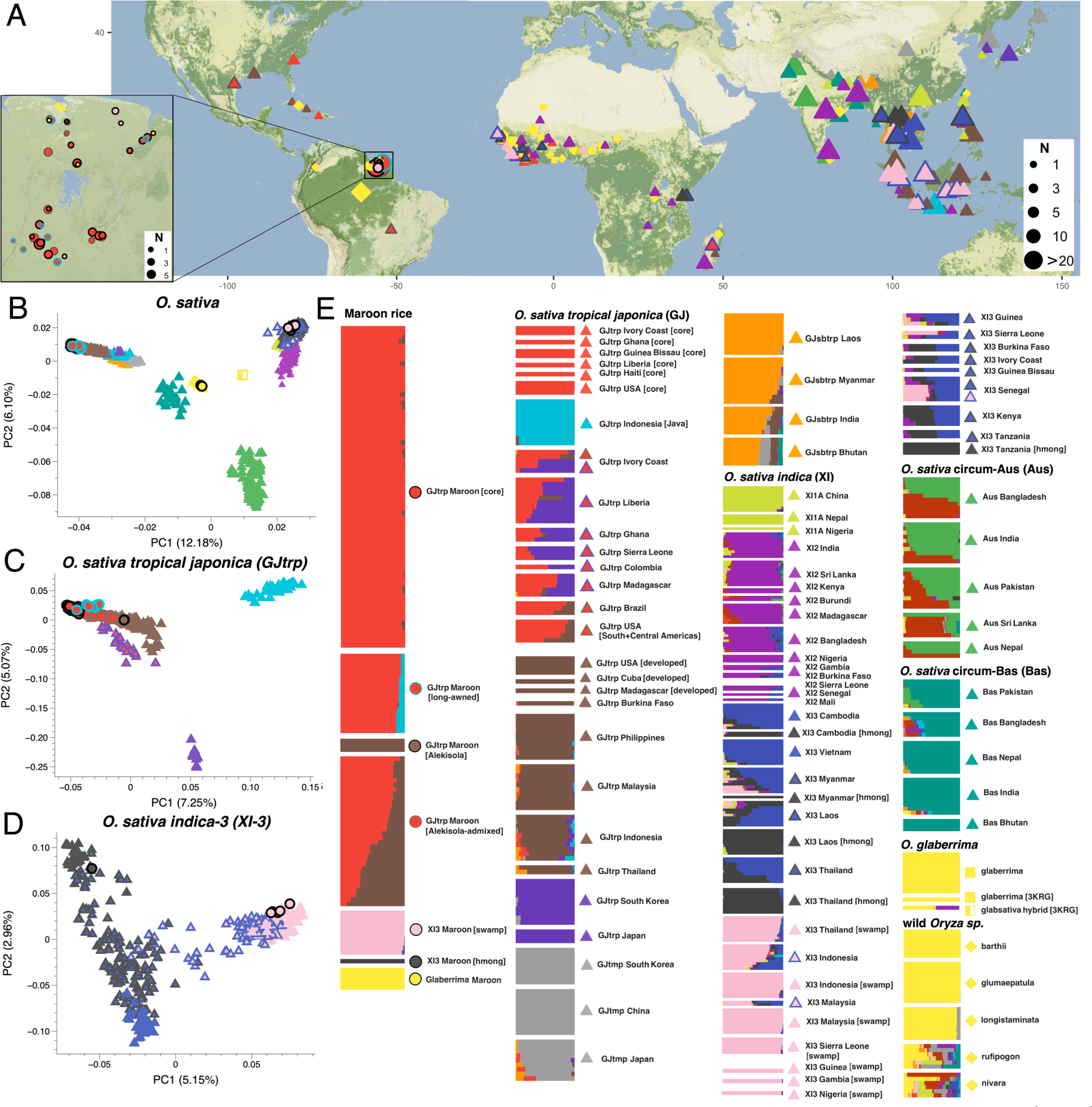
Global ancestry characterization of Maroon rice varieties in relation to worldwide rice diversity. Circles indicate Maroon varieties. Other symbols represent different rice species: O. sativa (triangles), O. glaberrima (squares) and wild rice (diamonds). Colors reflect the ancestry clusters and profiles assigned by ADMIXTURE at K=15. **(A)** Sampling locations of the rice varieties analyzed here, detailing the Maroon communities in the Guianas (box). Symbol size corresponds to sampling size (number of accessions). **(B)** PC space constructed from global O. sativa landrace diversity. The O. glaberrima and wild rice varieties are projected into the O. sativa PC space. PC1 separates O. sativa japonica varieties from indica varieties, and PC2 the circum-Aus varieties. O. glaberrima and wild rice varieties overlap poorly with O. sativa diversity and are pulled towards the center. (**C)** PC space constructed from global tropical japonica (GJtrp) varieties. The GJtrp Maroon core varieties cluster outside of the Southeast-Asian landrace diversity. The GJtrp Maroon long-awned varieties are shifted on a cline towards GJtrp Java, and the GJtrp Maroon Alekisola-admixed varieties plot between the GJtrp Maroon core and GJtrp Maroon Alekisola varieties. **(D)** PC space constructed from global O. sativa indica-3 (XI3) varieties. The first two PCs discern three distinct XI3 groups, and the XI-3 Maroon varieties cluster among two of them. **(E)** ADMIXTURE plot for K=15. A maximum of ten accessions was randomly selected per rice group, for the full data set with detailed individual labels, see Data S2. The ancestry profiles of the Maroon rice varieties cluster in seven genomic groups; four can be discerned among the O. sativa tropical japonica varieties, two among the O. sativa indica-3 varieties, and one non-sativa group that after additional ancestry characterization we identified as O. glaberrima, African rice.

### Seven rice genomic groups

Then we investigated how the Maroon varieties relate to the diversity of global rice landraces, and what genomic groups as defined in the 3K-RG project are represented (Wang, et al. 2018). For this we used Principal Component Analysis (Patterson, et al. 2006) (PCA, Figures 1B-D), and ADMIXTURE (Alexander, et al. 2009) for the unsupervised model-based clustering of global rice diversity (Figure 1E). Overall, we identified seven Maroon rice genomic groups. By far the majority (∼93%) of the Maroon varieties are *O. sativa* of the *tropical japonica* variety (‘GJtrp Maroon’, n=126), whereas *O. sativa indica-3* varieties account for a small subset (∼4%) (‘XI3 Maroon’, n=6) of the diversity (Figures 1B-D). A few Maroon landraces (∼3%) have ancestry that overlaps poorly with *O. sativa* diversity (Figure 1B). Maroons call these varieties *baaka alisi* (black rice) and *matu alisi* (forest rice), and they are morphologically similar to *O. glaberrima* (African rice). We hypothesize that these are Maroon varieties of *O. glaberrima* and labeled this group ‘Glaberrima Maroon’ (n=4).

Within the Maroon *O. sativa* diversity, the ADMIXTURE clustering model further distinguishes two *tropical japonica* and two *indica-3* Maroon rice groups with non-admixed ancestry profiles (Figure 1E and Data S2). The largest group consists of rainfed *tropical japonica* varieties with a large morphological diversity and traits characteristic of landraces, which we labeled ‘GJtrp Maroon core’ (n=73, including one previously published 3K-RG variety). The second group consists o*f tropical japonica* varieties referred to by the Maroons as *Alekisola*, which we labeled as ‘GJtrp Maroon Alekisola’ (n=3). The largest *indica-3* genomic group consists of wetland landraces that are nearly all grown by Okanisi Maroons in deep swamps in the Cottica region. We labeled these as ‘XI3 Maroon swamp’ (n=5). One *indica-3* variety is referred to by the Maroons as *Hmong*, most likely after the ethnolinguistic Hmong group from Laos and Vietnam, which we labeled ‘XI3 Maroon Hmong’ (n=1). Importantly, these Maroon rice genomic groups show distinct affinities to landraces in different world regions (Figure 1E and Data S2). Therefore, we hypothesize that these rice groups reflect various historical events during which rice was introduced to the Americas. Over time, several of these introduced rice varieties were incorporated into Maroon agriculture and grown and maintained as heirloom varieties by Maroon farmers with the seeds passed down from generation to generation.

In addition, the ADMIXTURE clustering model distinguishes two *tropical japonica* Maroon rice groups with dually admixed ancestry profiles (Figure 1E). The largest group contains varieties for which one ancestry component is found also in a non-admixed form in the ‘GJtrp Maroon core’ group and the other in the ‘GJtrp Maroon Alekisola’ group. We labeled this group ‘GJtrp Maroon Alekisola-admixed’ (n=35). The other admixed group consists of varieties with distinctively long, stiff awns, which we labeled ‘GJtrp Maroon long-awned’ (n=18, including two previously published 3K-RG varieties). These varieties carry one ancestry component that only appears in non-admixed form in *tropical japonica* landraces from Java. Since both admixed GJtrp Maroon groups have at least one ancestry component that is also found in non-admixed form among the Maroon varieties, we hypothesize that these rice groups resulted from local admixture. Maroon farmers may have discovered off-types that resulted from (spontaneous) crosses, and actively selected varieties with desirable traits.

### Rice from at least four historical periods

#### Transatlantic slave trade (1650-1825)

First, we confirmed our hypothesis that the four landraces that we labeled ‘Glaberrima Maroon’ are indeed Maroon varieties of African rice. *O. glaberrima* was domesticated from wild *O. barthii* ∼3,000 years ago along the Niger River (Meyer, et al. 2016), and except for Maroon communities (van Andel, van der Velden and Reijers 2016) is nowadays cultivated only in Africa. Around 1700*, O. glaberrima* likely was also cultivated in the Carolinas (USA) and elsewhere in the Americas, grown from seed stock that had been taken aboard of slave ships (Carney 2002). However, after the introduction of mechanical milling devices, *O. sativa* replaced *O. glaberrima* as an export crop. In genomic distance analyses based on shared-IBS, the ‘Glaberrima Maroon’ varieties form a cluster with a previously published *O. glaberrima* variety obtained from Maroons in Suriname (van Andel, Meyer, et al. 2016), and fall within a larger cluster with West African *O. glaberrima* varieties (Figure S3). Although outgroup *f_3_*-statistics cannot conclusively discern whether the Maroon *baaka alisi* and *matu alisi* varieties share more ancestry with *O. glaberrima* varieties or *O. barthii* (Figure S4), the results of various *f_4_*-statistics indicate that the Maroon varieties form a clade with *O. glaberrima* varieties from West Africa to the exclusion of *O. barthii* (Table S1 and S2). These findings are congruent with previous genomic studies that showed that rice landraces with similar morphological characteristics and Maroon nomenclature are *O. glaberrima* varieties that originated from West Africa and were introduced during the slave trade to the Guianas (van Andel, Meyer, et al. 2016; Veltman, et al. 2019).

Second, we aimed to characterize the ancestry of the morphologically diverse *tropical japonica* landraces in the ‘GJtrp Maroon core’ group (n=73). In ADMIXTURE, the ancestry component for this group is also found maximized in a few landraces from West Africa (Guinea Bissau, Ghana, Ivory Coast, and Liberia), one from Haiti and four early-registered cultivars from the USA (Figure 1E and Data S2). Intensive trade networks connected these areas during colonial times, starting in the early 16^th^ century (Eltis and Richardson 1995). *(Tropical) japonica* varieties from the Philippines and Malaysia were introduced by the Portuguese to West Africa around 1520 (Nawani 2013; Gilbert 2015), and shipped from West Africa together with enslaved Africans to Brazil from the 1530s onwards (Carney 1998). As such, the similar ancestry found in the landraces of the ‘GJtrp Maroon core’ group and varieties in other regions of the Americas and West Africa may reflect a relict genomic signature that traces back to transatlantic trade that connected these areas during early colonial times. To investigate this in more detail, we combined individual-level genomic distance analyses with various population-level symmetry *f_4_*-statistics. We found that the landraces in the ‘GJtrp Maroon core’ group show the highest levels of shared genetic drift with ‘core’ landraces in West Africa, Haiti and some of the early-registered USA cultivars, and lower levels with other varieties from the Americas (e.g., Brazil, Colombia), (West) Africa and Asia (Figure 2A and S5, Data S3). These findings hence underline a direct genomic link between the ‘GJtrp Maroon core’ group and other ‘core’ varieties from the Americas to the ‘core’ landraces in West Africa.

**Figure 2.**
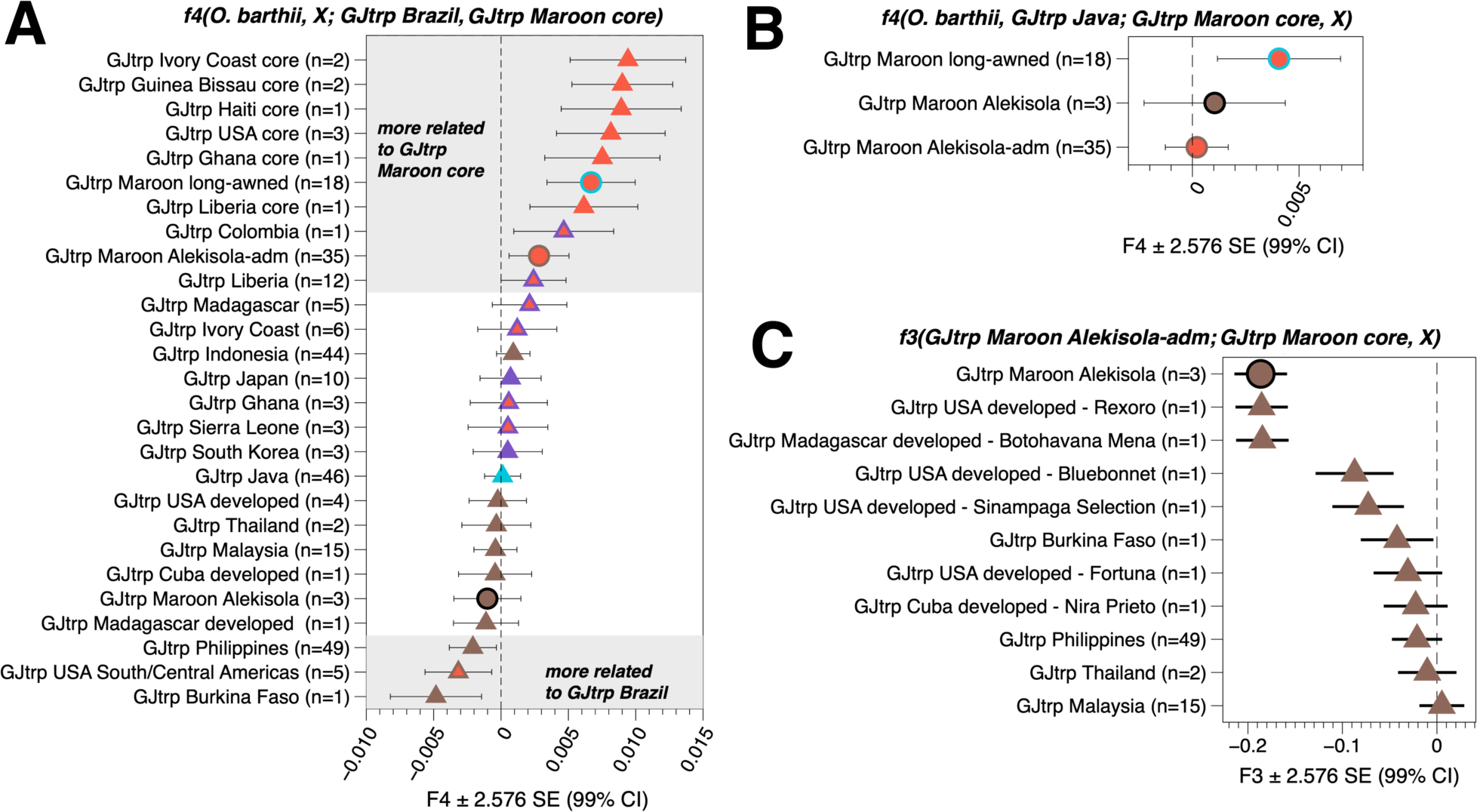
F-statistics for the ancestry characterization of the various tropical japonica (GJtrp) Maroon rice groups to trace their geographical origins. Error bars reflect 2.576 SEs, corresponding to 99% confidence intervals (CIs). **(A)** F_4_-statistics that compare the ancestry of the ‘GJtrp Maroon core’ and ‘GJtrp Brazil’ varieties to measure their differences in shared genetic drift to other GJtrp groups (X). The ‘GJtrp Maroon core’ group significantly shares more alleles with other ‘core’ subsets in the Americas and West Africa, indicating a close genomic link. **(B)** Admixture f_4_-statistics that test for directional admixture of ‘GJtrp Java’ ancestry in the various ‘GJtrp Maroon’ groups. ‘GJtrp Maroon core’ represents the ancestry of the progenitor population, i.e., the tropical japonica diversity cultivated by the Maroons prior to admixture. Only the ‘Maroon GJtrp long-awned’ group shows a significantly positive f_4_-statistic (Z = 3.625), indicating that the Java ancestry is specific to this group. **(C)** Admixture f_3_-statistics for the ‘GJtrp Maroon Alekisola-admixed’ group. The statistics test what combination of ancestry from the ‘GJtrp Maroon core’ group and another GJtrp group X approximates the gene pool of the ‘GJtrp Maroon Alekisola-admixed’ group best. The most extreme negative f_3_-values indicate the strongest admixture signals and are found for the local ‘GJtrp Maroon Alekisola’ varieties (99% CI: −0.214 < F_3_ < −0.159) and the closely related early developed cultivars Rexoro and Botohavana Mena. The admixture f_3_-values are significantly less negative for more distantly related landraces from the Philippines (99% CI: −0.048 < F_3_ < −0.005) and other regions in Southeast Asia.

Maroon oral accounts and colonial archival records provide evidence of rice imports from West Africa to the Guianas to feed soldiers and enslaved Africans on the plantations (Fleury 1994; Carney 2005; van Andel 2010). The genomic analyses confirm that the Maroon farmers have preserved a part of this rice diversity, including both *O. sativa tropical japonica* and *O. glaberrima* varieties. Among the *tropical japonica* varieties that fall in the Maroon ‘core’ group are varieties named *Ma Paanza*, *Alisi Seei*, *Baakapautjaka and Agbosó* (Data S2 and S3). According to Maroon oral history, these are old varieties that were named after specific female ancestors or early runaway groups that brought rice to the Maroon villages (Pinas, et al. 2023). These old varieties in the ‘GJtrp Maroon core’ group and the *O. glaberrima* varieties typically have a short maturation period of circa three to four months (van Andel 2010; Van Andel, et al. 2019). According to the Maroons, these were particularly suitable crops for (re)settlement after escape from plantations and military expeditions to recapture Maroons (van Andel 2010).

#### Javanese indentured laborers (1890-1930)

Subsequently, we characterized the ancestry of the *tropical japonica* varieties in the ‘GJtrp Maroon long-awned’ group (n=18). The major ancestry component is shared with the ‘GJtrp Maroon core’ varieties. However, ADMIXTURE distinguishes a minor ancestry component (5.9-13.3%) that is characteristic of *tropical japonica* landraces from Java (Figure 1E and Data S1). Since many Javanese *tropical japonica* have similar long and stiff awns (Iskandar and Ellen 1999) this raises the possibility that this trait admixed into some of the Maroon rice varieties. Indeed, using various admixture *f_4_*-statistics we found a significant admixture signal for ancestry from Javanese landraces only in the ‘GJtrp Maroon long-awned’ group (Z = 3.625) and not in other (Maroon) landrace groups (Figure 2B and S6A). Moreover, a significant admixture signal is obtained only for landraces from Java, and not for sources from other geographical regions (Figure S6B). The characteristic Java ancestry led us to link the origin of the Maroon long-awned varieties to a specific historical context. After the abolishment of slavery in 1863, the Dutch tried to keep their plantation economy in Suriname viable by contracting laborers from India and Java (Hoefte 1998). After their contract ended, the majority of the Asian workers settled in Suriname as smallholders and continued cultivating rice for subsistence (Maat and van Andel 2018). Until the 1980s, Javanese in Suriname grew their traditional rice varieties in food plots on the few remaining coffee and sugar plantations where they were employed. Young Maroon men, searching for wage labor in coastal Suriname, often ended up working on these plantations as well (Price 1993). Here, they probably obtained long-awned Javanese rice and gave the planting stock to their families in the interior. Notably, although laborers from India also brought rice with them to Suriname (Maat and van Andel 2018), we did not (yet) find an unambiguous genomic link between the Maroon rice varieties in our data set and landraces in India.

#### USA cultivars (1936 onwards)

We then aimed to further characterize the ancestry of the *tropical japonica* varieties that the Maroons refer to as ‘*Alekisola’*, which we hence labeled as ‘GJtrp Maroon Alekisola’ (n=3). These are upland varieties with elongated, awnless grains and smooth husks, and a circa five-month growth period, which is longer than that of the landraces in the ‘GJtrp Maroon core’ group. In the ADMIXTURE clustering analysis a similar ancestry profile is found for landraces in Southeast Asia, predominantly the Philippines, and some early-developed USA cultivars, including the glabrous-hulled *Rexoro* that was developed in 1926 in Louisiana (Rutger and Mackill 2008) (Figure 1E, Data S2). *Rexoro* quickly became one of the most popular and economically lucrative rice varieties of that time and was shipped to various places over the world for trade and breeding experiments (Lu, et al. 2005). Historical literature indicates that *Rexoro* was introduced to the Maroons in Suriname in 1936 (Stahel 1944). We hence investigated whether the Maroon rice varieties named ‘*Alekisola’* are local variants of *Rexoro* and/or other early-registered USA cultivars, or more related to landraces in the Philippines. Outgroup *f_3_*-statistics indicate that the level of shared genetic drift between the ‘GJtrp Maroon Alekisola’ varieties and the USA early-developed cultivar *Rexoro* is significantly higher than for ‘GJtrp Philippines’ (Figure S7). Similarly, *Alekisola* varieties cluster with USA *Rexoro* and show a very high level of IBS-sharing (Data S3). These results indicate that the *Alekisola* and *Rexoro* varieties are closely related and underline the historical evidence that *Rexoro* was sent to the Maroons (Stahel 1944; Maat and van Andel 2018).

In addition, we find that a substantial subset of the Maroon rice diversity, which we labeled ‘GJtrp Maroon Alekisola-admixed’ (n=35), appears dual admixed between varieties that fall in the ‘GJtrp Maroon core’ and ‘GJtrp Maroon Alekisola’ groups with the *Alekisola* component seemingly accounting for up to ∼90% in ADMIXTURE (Figure 1E, Data S2). Many varieties in this group have names like *Alekisola*, *Lexola,* and *Sola,* and are especially popular among Saamaka rice farmers, who distinguish several types by the color of their husk (Pinas, et al. 2023). However, *tropical japonica* landraces from Brazil, as well as five early-registered USA cultivars that were collected as landraces in Central and South America, and a few landraces from West Africa, similarly show a dual admixed ancestry profile (Figure 1E, Data S2). Therefore, we investigated whether the varieties in the ‘GJtrp Maroon Alekisola-admixed’ group more likely resulted from Maroon crossings between varieties in the local ‘GJtrp Maroon core’ and ‘Alekisola’ groups, or were introduced from places outside the Guianas, such as Brazil. Admixture *f_4_-*statistics, taking either ‘GJtrp Brazil’ or ‘GJtrp Maroon core’ as a progenitor source, show that the ‘GJtrp Maroon Alekisola-admixed’ group has significant additional ancestry that best matches the ancestry of ‘GJtrp Maroon Alekisola’ or the closely related ‘GJtrp USA early-developed’ cultivars (e.g., *Rexoro*) (Figure S8). The local ‘GJtrp Maroon core’ landraces are a closer progenitor source than those from Brazil, since for the statistic *f_4_(O. barthii, GJtrp Maroon Alekisola-admixed; GJtrp Brazil, GJtrp Maroon core)* the ‘GJtrp Maroon Alekisola-admixed’ and ‘GJtrp Maroon core’ group show a significant excess of shared genetic drift (Z = 3.27) (Figure 2A). The differential ancestry between ‘GJtrp Brazil’ and ‘GJtrp Maroon core’ group that is driving this statistic is associated with ancestry from ‘GJtrp Philippines’ (Z = −3.08), and not ‘GJtrp Maroon Alekisola’ (Z = −1.05) or the USA early-registered cultivars (Z = −0.29) (Figure 2A). Subsequently, we tested the hypothesis that the ‘GJtrp Maroon Alekisola-admixed’ gene pool resulted from local admixture more directly, using conservative admixture *f_3_*-statistics (Figure 2C and S9). Indeed, the strongest signals for admixture are found for the local ‘GJtrp Maroon *Alekisola*’ varieties (or the closely related early-developed *Rexoro* cultivar from the USA) (Figure 2C), and ‘GJtrp (Maroon) core’ varieties (Figure S9). This indicates that the ‘GJtrp Maroon Alekisola-admixed’ group on average can be considered a result of local admixture between varieties in the ‘GJtrp Maroon core’ and ‘GJtrp Maroon Alekisola’ groups. We note, however, that the ‘GJtrp Maroon Alekisola-admixed’ group appears to have an additional genomic substructure. The varieties show differential affinities to *Alekisola* and USA early-developed cultivars on one hand, and to landraces from South America and a few from West Africa that share relatively more ancestry with landraces from the Philippines, on the other (Data S3). In comparison, the underlying substructure in West African varieties appears to be predominantly associated with ancestry characteristic for *tropical japonica* landraces in Japan and South Korea (Figure 1E and Table S3). Additional genomic data for (historical) rice landraces from South/Central America and West Africa will be crucial for elucidating the genomic affinities underlying the additional substructure of the *tropical japonica* landraces on both sides of the Atlantic in more detail.

#### Creoles in coastal Suriname (after 1850?)

In addition to the predominant *tropical japonica* varieties, some Maroon farmers have also preserved two distinct subsets of *indica-3* landraces (Figures 1D and E). The largest genomic group ‘XI3 Maroon swamp’ (n=5) is cultivated almost exclusively by Okanisi Maroons in swamps in the Cottica region (Figure S1). These wetland landraces have very tall stems (> 2 m), often a red pericarp, and a long growth season of circa seven months. In ADMIXTURE, landraces that are grown in deep mangrove swamps in Southeast Asia (Indonesia, Malaysia, Thailand) and West Africa (Sierra Leone, Guinea, Nigeria, Gambia) show a similar ancestry profile to the ‘XI3 Maroon swamp’ varieties (Figure 1E and Data S2). Hence, we investigated whether the ‘XI3 Maroon swamp’ varieties are more related to varieties in Southeast Asia or West Africa. We find that the ‘XI3 Maroon swamp’ varieties cluster as a group among swamp landraces from Indonesia, in particular Java, Sumatra, and Sulawesi, and not among West African varieties (Data S4). Congruently, symmetry *f_4_-*statistics indicate that the ‘Maroon XI3 swamp’ varieties are more closely related to XI3 landraces in Indonesia, whereas those from West Africa are more closely related to XI3 varieties from Malaysia and Thailand (Figure S10). This makes it unlikely that the Maroon varieties derived directly from varieties in West Africa. The historical details about how the Maroons obtained these varieties are difficult to establish. Some Maroon farmers said they received this rice from Creoles, descendants of enslaved Africans who became smallholder farmers after emancipation and were addressed respectfully as *masaa* (master) (Pinas, et al. 2023). Congruently, historical records attest that a group of free Okanisi returned from the interior of Suriname to the coastal Cottica swamps around 1850 (Thoden van Velzen 2022). Other Okanisi said they received this rice from city Creoles with a high social position, who are also addressed as *kioo* (Creole) or *masaa*, and visited Maroon villages as teachers, missionaries, and agricultural extension officers in the period 1930-1980. Hence, more detailed reconstructions regarding how, when, and how often the Maroons obtained these swamp varieties require further genomic, ethnobotanical, and archival investigation.

#### Hmong refugees in the Guianas (1991)

Lastly, we investigated the genomic affinities of a distinct Maroon *indica-3* upland rice variety with elongated grains, a white husk, short white hairs, and referred to by the Maroons as *Hmong* rice (‘XI3 Maroon Hmong’). The Maroons explained that they obtained this *indica-3* variety from Hmong refugees (Pinas, et al. 2023). The Hmong are an ethnolinguistic group from the northern mountains of Laos, Vietnam, and Thailand. During the Vietnam War, many Hmong fled their native area and some settled in French Guiana around 1977 (Clarkin 2005). In ADMIXTURE, the Maroon *Hmong* variety is maximized for an ancestry component that is predominantly found in *indica-3* landraces in Laos and Thailand (Figure 1E and Data S2). With individual-level genomic distance analyses, we found a close genomic affinity to broad subset of *indica-3* landraces from Thailand, Laos and Cambodia, and a very close genomic affinity to one variety from Thailand, one from Myanmar and three from Tanzania (Figure S11 and Data S4). Since we could not find historical records that mention that the Hmong people migrated to Tanzania, we suspect that these varieties were exchanged through rice breeding stations. As a result of the civil war in Suriname (1986-1992), large numbers of Okanisi Maroons from the Cottica found refuge in French Guiana. Although the French authorities did not allow them to farm, some Maroons worked illegally on the Hmong farms in 1991 (Hoogbergen and Polimé 2002). We posit that the Maroon *Hmong* variety was most likely obtained directly from the Hmong during this time when farmers from both groups met in French Guiana. After the civil war, several Maroons returned to the Cottica. There they continued to grow this *Hmong* variety as a welcome addition to their rice portfolio, which had been severely diminished by the civil war and forced displacement.

## Discussion

By combining genomic insights on the likely geographical origin of the newly sequenced 136 Maroon rice landraces with information from the Maroon community and historical records, we linked the origin of the Maroon rice to at least four different historical events (Figure 3A). We find that the largest subset traces back to the transatlantic slave trade from West Africa, while other rice varieties were obtained from interactions after emancipation, linked to Java, the USA and Vietnam. These findings underline that Maroon farmers are capable seed selectors, for conservation of appreciated varieties and for innovating their varietal stock.

**Figure 3.**
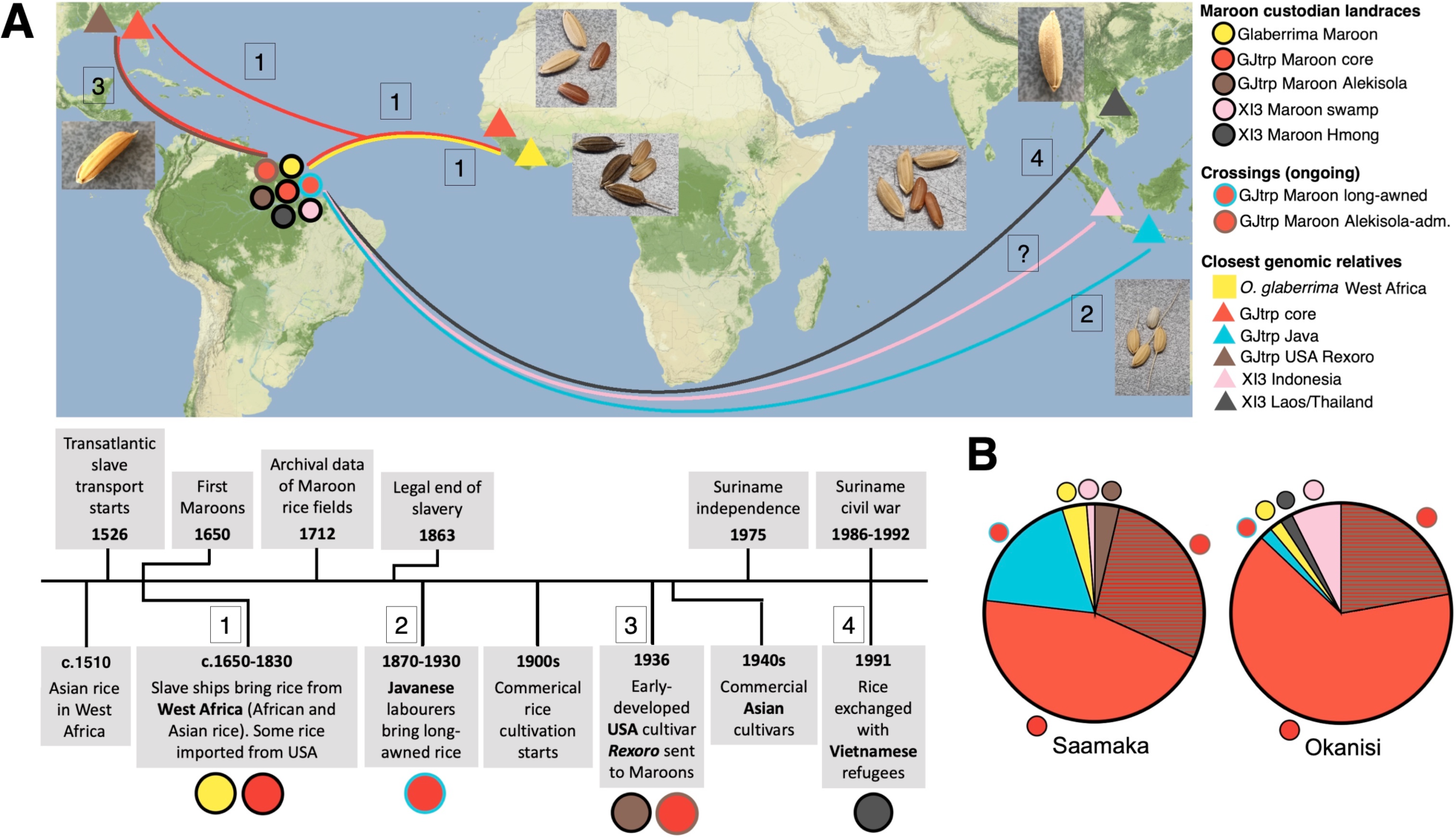
Maroon farmers have preserved rice varieties from at least four introduction events to the Guianas during (post-)colonial history. **(A)** Summary map and timeline showing the geographical connections and chronology of the historical contexts associated with the Maroon rice genomic groups. [1]: Tropical japonica and African ‘black’ rice varieties from West Africa during the transatlantic slave trade (1650-1825). [2]: Long-awned tropical japonica from indentured laborers from Java (1890-1980). The Javanese ancestry is only detected in admixed form in the Maroon tropical japonica varieties. [3]: Tropical japonica USA cultivar Rexoro (1936 onwards). [4]: Indica-3 variety from Hmong refugees (1991). [?]: Cottica Maroons receive swamp landraces that ultimately come from Indonesia. **(B)** Relative proportions of the various Maroon rice genomic groups observed among Saamaka and Okanisi farmers. Despite their preference difference for certain rice traits, farmers of both Maroon groups cultivate varieties from nearly all genomic groups, demonstrating a shared motivation for maintaining crop diversity.

Moreover, our genomic analyses identified types of rice never documented before, i.e., long-awned *tropical japonica* varieties with Java ancestry and some of the Alekisola-admixed *tropical japonicas*. This shows that Maroon farmers are not only avid collectors but also active selectors of new rice varieties, derived from intentional and/or spontaneous crosses, to further increase their stock diversity. We posit that the high landrace diversity currently grown by Maroons reflects the farming expertise they inherited from their (African) ancestors (Mokuwa, et al. 2013). Ignored by the agricultural modernization initiatives of the colonial government and the independent Suriname state, the Maroons have followed their own trajectory of landrace conservation and agricultural improvement.

A major cultural driver for maintaining a high rice diversity in the Maroon farming system is the exchange of seeds of successful and desired varieties. This mostly takes place between families, neighbors, and friends within the community. Village ceremonies, such as funerals, are typical events where gifts and tokens are exchanged, including seeds (Van Andel, et al. 2019; Pinas, et al. 2023). Although Maroon groups have distinct preferences for certain rice traits, e.g., long awns among the Saamaka and swamp rice among the Cottica Okanisi (Pinas, et al. 2023), their seed exchange and cultivation strategies are similarly focused on creating and maintaining rice diversity (Figure 3B). Importantly, among both the Saamaka and Okanisi communities the newer *Alekisola*(-admixed) and *indica* varieties never fully replaced the older *tropical japonica* landraces that represent the Maroon communities’ roots to West Africa. Taken together, the results of our genomic analyses are congruent with many events as preserved in Maroon oral history and may further strengthen their cultural identity.

Moreover, this study underlines the central importance that the Maroons had in the development of rice farming practices in the Americas. Although rice was purchased as bulk food by slave traders in West Africa and grown for subsistence on plantation food plots on both sides of the Atlantic (Wood 1975; Carney 1998; Dragtenstein 2017) (Supplementary Material: Evidence #1-4), colonial authorities frequently overstated their involvement in the discovery and development of productive rice varieties and farming innovations that were economically lucrative (Carney 2002). Early archival documents on the crop in Suriname and Brazil just mention “rice that grows here” (Supplementary Material: Evidence #1-2) and remain silent on those who found the seeds and knew how to plant and harvest them. Enslaved Africans were often allowed to grow their own food on small provision fields (Fleury 1994; van Andel, van der Velden and Reijers 2016) and may have planted rice in secret as an ‘anti-commodity’ crop (Maat 2015), hidden between other crops as an act of protest against the slave-based plantation system and in memory of their African heritage (Carney 2005; Maat and van Andel 2018). After their escape to freedom and once safely settled in the forested interior, Maroons were free to invest time and space into the farming systems of their choice, combining familiar African food crops (e.g., rice, okra, plantains) with crops of indigenous origin like cassava, tobacco, and sweet potato (Price 1991; van Andel, van der Velden and Reijers 2016). The fact that colonial authorities in the Guianas have sent out military raids for almost two centuries to destroy Maroon provision fields (Dragtenstein, et al. 2002) (Supplementary Material: Evidence #5) attests to the vast knowledge of Maroon farmers to establish highly productive farming systems in different ecosystems.

As such, this study provides a case study that demonstrates the power of cross-disciplinary crop genomics research to obtain a more comprehensive perspective on a time of the human past for which the historical record may be biased or incomplete. Similar studies can be envisioned for landraces of other ‘cultural heritage’ crops of (Indigenous) communities who may have preserved their history on their own farms, to both reconstruct and honor the past.

## Materials and Methods

### Ethnobotanical surveys

Fieldwork on Maroon rice farming was carried out in French Guiana and Suriname in 2017, 2021, and 2022. Before the start of each fieldwork period, we obtained permission from the traditional Maroon authorities and prior informed consent from each individual rice farmer. In addition, we obtained consent and permission of the Suriname National Rice Research Institute (SNRI/ADRON). We interviewed 88 Maroon rice farmers (21 in 2017, 67 in 2021-2022, almost all women), in Dutch, French, or one of the Maroon languages. We asked questions about the rice varieties grown, their local names, the meaning of these names, specific varietal traits, farmers’ motivation for rice cultivation, specific rituals involving rice, and oral history associated with rice and its origins (Van Andel, et al. 2019; Pinas, et al. 2023; van Andel, et al. 2023).

### Rice samples

Our primary focus was to collect as much Maroon rice diversity as possible, rather than to obtain an accurate account of the diversity preserved by individual farmers. Hence, when we encountered a rice variety with morphological characteristics and Maroon rice name that was identical to that of a variety that we had already collected, we did not sample it again. Herbarium vouchers of rice plants were collected using standard botanical methods. Duplicates were deposited at the Herbier IRD du Guyane (CAY) in Cayenne (French Guiana), the National Herbarium of Suriname (BBS) in Paramaribo, and at Naturalis Biodiversity Center (L) in Leiden, the Netherlands. Living seeds of most varieties have been deposited with the associated data on locality, varietal information, and farmer’s name at the Suriname National Rice Institute (SNRI/ADRON) at Nickerie for further propagation, phenotyping, and germplasm storage. Dead seed samples were stored at the Naturalis Economic Botany Collection. For each collected variety, we reported morphological and agronomical characteristics.

### Historical data collection

The digital archives of the Dutch West India Company and the Society of Suriname, both housed by the National Archives in the Hague, the Netherlands, were queried for the search terms ‘rice’, ‘rijst’, ‘rys(t)’, ‘reis’, ‘kost’ (provision), ‘gronden’ (grounds). A literature survey was carried out on rice cultivation in the Guianas and Dutch Brazil during the early colonial period, and archival records mentioned in these studies were verified.

### Sequencing data

DNA extracts and libraries for the rice specimens were generated from leaf, stem, or seed material from living rice plants that were either collected in the farmer’s field or fresh leaf from germinated seeds in the Netherlands (Data S1A). Preferably, we used leaf material when available. The tissue samples were dried with silica gel beads and stored in dark containers for conservation. The tissue samples were ground using the Retsch Mixer Mill MM400 while frozen in liquid nitrogen. For each rice specimen, a DNA extract was generated from typically 10-20mg powder with a standard CTAB-based method (https://openwetware.org/wiki/Maloof_Lab: 96well_CTAB). We added 2% PVP-40 to the CTAB buffer to remove phenols and polysaccharides. DNA yields in the extracts were 12-140 ng μl^-1^. Then 5uL of (normalized) DNA extract was converted into double-stranded DNA libraries with unique dual indexes (Data S1A) using tagmentation with an in-house protocol for the Illumina DNA Prep kit (formerly Nextera DNA Flex). Our in-house protocol is based on a previously published ‘hack’ protocol (Gaio, et al. 2022), but we added normalization steps before and after tagmentation to balance the DNA concentration, hence raw sequence read output, among the rice accessions. The ‘hack’ protocol allows us to use the Illumina kit for 1450 instead of 24 library preps. Sequencing was done on the Illumina Novoseq6000 platform with 2×150bp paired-end read configuration to a target genome-wide sequencing depth of ∼14X at the Core Facility of Novogene Europe in Cambridge or GenomeScan in Leiden. The sequenced reads were demultiplexed according to the expected index pair for each library, allowing one mismatch per 8 bp index. Sequencing data for these accessions are available from the SRA under Bioproject accession number PRJNA1019116.

### Read processing, quality control and genotyping

The grenepipe v0.7.0 snakemake pipeline was used for read processing, quality control and calling SNPs (Czech and Exposito-Alonso 2022) (https://github.com/moiexpositoalonsolab/grenepipe). Prior to starting the read processing pipeline, the fastq format files from different sequencing runs for the same library were merged. We used *fastp* v0.20.0 (Chen, et al. 2018) to clip adapters, Ns stretches and PolyG tails of the reads. We merged paired end reads into a single sequence for regions with a minimal overlap of 30bp, and only kept merged sequences ≥55bp. Sequence reads were mapped against the Nipponbare IRGSP-1.0 reference genome (Kawahara, et al. 2013), a *temperate japonica* rice cultivar, using the global aligner BWA v.0.7.17 in ‘mem2’ mode (Vasimuddin, et al. 2019) producing .bam files. The mapped reads were sorted, indexed, and filtered (mapQ 60) using samtools v1.3 (Li, et al. 2009). Duplicate reads, identified by comparing sequences in the 5’ positions of both reads and read pairs, were removed with MarkDuplicates in PICARD v2.23.0 (http://broadinstitute.github.io/picard/). The .bam files were used to call diploid haplotypes across all rice accessions with Freebayes v1.3.1 (Garrison and Marth 2012), parallelized for contigs (e.g. chromosomes), while forcing allele calls at 29 million known SNP positions (https://snp-seek.irri.org/_download.zul) using the option *--haplotype-basis-alleles* combined with BEDTOOLS v2.30 (Quinlan and Hall 2010) for region intersection (bedtools *intersect* -a). These 29 million SNP positions were previously validated as segregating among >3000 rice accessions of the 3K-RG project for reads mapped to the Nipponbare IRGSP-1.0 reference genome, and include the 404k core set. The VCF files for each contig were concatenated, indexed and compressed with PICARD MergeVcfs, resulting in a single VCF file with raw genotype data for a set of 29M genome-wide SNPs across all Maroon rice accessions. From this raw set of SNPs we removed all non-biallelic sites. Subsequently, with GATK v4.1.4.1 (McKenna, et al. 2010) we subjected it to a series of hard filtering steps (--filter-name snv-hard-filter) to remove SNP sites with low-quality genotypes normalized by depth (QD < 2.0), mapping quality (MQ < 60.0 and MQRankSum < −12.5), read position bias from Wilcoxon’s test (ReadPosRankSum −8.0) and strand bias from Fisher’s test (FS > 60.0). Using an in-house python script we extracted the genotypes for all SNP positions in the 3K-RG 404k core SNP panel (https://snp-seek.irri.org/_download.zul), and transformed the VCF file into packedped files with PLINK v1.90b (Purcell, et al. 2007; Chang, et al. 2015), and subsequently into eigenstrat files using *convertf* from the EIGENSOFT package v7.2.0 (https://github.com/DReichLab/EIG).

### Comparative dataset

We made a core comparative data set for global rice landrace diversity that we based on the 3K-RG 404k CoreSNP Dataset in PLINK format provided by the Toronto International Data Release Workshop Authors, available via https://snp-seek.irri.org/_download.zul. The 404,388 bi-allelic SNPs in the 3K-RG 404k CoreSNP panel were selected to optimally discern between the major rice genomic groups of Asian (*O. sativa*) rice, as well as discern some (eco-)geographical structure among rice varieties within each of the major rice genomic groups. From the 3,024 rice accessions in this 3K-RG 404k CoreSNP Dataset we selected the rice landraces from Asia (n=1,064) for which landrace status was previously validated by Gutaker *et al*. (Gutaker, et al. 2020). In addition, using a similar approach to Gutaker *et al*., we selected 3K-RG rice accessions from Africa (n=97) and the Americas (n=11) that in Genesys (https://www.genesys-pgr.org) were annotated with ‘traditional’ status or as an inbred lineage descendent thereof, to validate them as putative landraces in downstream analyses. To expand the comparative data set further we downloaded sequencing data for USA early-registered cultivars (Vaughn, et al. 2021) (n=12 [SRA project accession PRJNA603026]), *O. glaberrima* varieties from Africa (Meyer, et al. 2016) (n=20 [PRJNA315063, PRJDB4713]) and from Suriname (van Andel, Meyer, et al. 2016) (n=1 [PRJNA514989]). Moreover, we downloaded sequence data for representative varieties for diploid wild *Oryz*a species (AA genomes) found in the Americas (*O. glumaepatula*: n=20 [PRJDB4703], *O. longistaminata*: n=9 [PRJDB4705]), Africa (*O. barthii*: n=8 [PRJNA30379]), and Asia (*O. nivara:* n=6 [PRJNA202926]), *O. rufipogon* (Huang, et al. 2012): n=6 [PRJEB2829]). The sequencing data was downloaded in fastq format from the Short Read Archive (SRA) using the FASTQ-DUMP tool v.2.8.2 with the option to split reads into forward, reverse and trimmed. Then the reads were processed and genotyped following the same pipeline as the newly sequenced Maroon rice varieties. The resulting 404k SNP genotype datasets were merged with that for the Maroon rice, and subsequently with that of the (candidate) landraces extracted from 3K-RG 404k CoreSNP Dataset.

#### SNP filtering

From the ‘404k core dataset’ set, we filtered out SNP markers with PLINK v.1.90b (Purcell, et al. 2007; Chang, et al. 2015) that were determined as outliers with the 1.5xIQR rule (more extreme than the 3th Quartile + 1.5 * IQR) regarding genotype missingness (Fmiss) and heterozygosity ratio (HetRateObs). The latter is to remove SNPs that we expect are the result from spuriously mapped reads that are from tandem duplicated genomic regions (Gutaker, et al. 2020). This filtering step were applied to the full dataset, as well as separately for 1) Maroon landraces 2) *indica* landraces, 3) *japonica* landraces, 4) circum-aus landraces, and 5) *O. glaberrima*/wild *Oryza* species. Genotype missing rates for markers (Fmiss) were determined with *--missing*, and SNPs with excess rates (typically Fmiss > 0.25) were removed from the dataset with *--geno.* To determine the threshold for an excess heterozygosity ratio, we first calculated the genotype counts per SNP with PLINK using *--freqx.* From the data in the .freqx file we subsequently calculated the heterozygosity ratio per SNP, here defined as: HetRateObs = C(Het) / (C(Hom A1) + C(Het) + C(Hom A2)). This corresponds to the heterozygosity count divided by the total genotype count (not including missing genotype counts). The IDs of SNP markers with outlier heterozygosity rates (typically > 0.20) were collected in a bad_snp_list, and subsequently removed from the dataset with *--exclude*. Then the SNP set was subjected to a two-step LD pruning procedure. The first step was carried out with the INDEP-PAIRWISE function in windows of 10 kb with variant shift = 1 and r2 = 0.8. The second step was carried out with the same function in windows of 50 variants. These filtering steps resulted in 109,203 highly informative SNPs (‘110k panel’). In contrast to the 404k core panel (Figure S2A), the calling rates for non-missing genotypes in the 110k panel are similar for the Maroon (95.0 ± 4.3%) and 3K-RG rice landraces (97.0 ± 4.3%) (Figure S2B). The average number of reads supporting a SNP genotype call (genotype depth) for the Maroon varieties is 10.3 ± 6.1X (mean ± s.d.), which overlaps with the 14.4 ± 6.9X mean genotype depth for the 3K-RG project accessions (Wang, et al. 2018) (Figure S2C). We caution that, similar to the 404k core panel, also the 110k core panel may not be as effective in resolving genomic affinities with non-sativa rice species, because a relatively large fraction of the SNPs is non-segregating.

#### Individual filtering

We removed rice varieties from the dataset with high marker missingness, using --*remove.* Similar to the SNP filtering steps, thresholds for outliers were based on the 1.5xIQR rule and determined separately for the different subsets of data. In addition, we filtered out landraces with mixed ancestry based on their joint genetic clustering in PCA and ADMIXTURE. These in general include, but were not limited to, 3K-RG varieties labeled as ‘admix’, ‘GJadm’ or ‘XIadm’, as well as some of the 3K-RG rice varieties from Africa and the Americas that we had preselected as ‘candidate landraces’.

### Principal Component Analyses (PCA)

PCA analyses were run with the program *smartpca* from the EIGENSOFT package v7.2.0 (https://github.com/DReichLab/EIG) on the global diversity of *O. sativa* landraces, as well as on the subsets of *tropical japonica* and *indica-*3 landrace diversity. In the PCA for global *O. sativa* diversity the *O. glaberrima* and wild *Oryza* varieties were projected on the PCs with the option *‘lsqproject: YES.* The variance explained by each principal component (PC) was calculated as the dimension’s eigenvalue divided by the sum of all positive eigenvalues, and the first two PCs were plotted.

### Population assignment

We conducted an unsupervised genomic clustering of global rice diversity using ADMIXTURE v.1.3.0 (Alexander, et al. 2009). To reduce the impact of different inbreeding levels among the rice varieties on the clustering results we reduced diploid to haploid calls by randomly sampling one allele. We ran ADMIXTURE with fivefold cross-validation varying the number of ancestral populations between K = 2 and K = 20 in 100 bootstraps with different random seeds (Data S1C, Data S2). The model did not converge to a global optimum (lowest cross-validation error) and continued to marginally improve with increasing *k*-means because of further breakdowns in geographical structure (Figure S12). This results from the rationale behind the design of the SNP panel to capture a high proportion of common variance *within* each of the 3K-RG major genomic groups to discern additional geographical structure. As such, the ADMIXTURE model starts to discern broad ecotypes within indica-3 (from K=7) and *tropical japonica* (from K=8) varieties before it has discerned the nine major rice genomic groups as defined in the 3K-RG project (at K=12). From K=16 the breakdown into increasingly smaller ecotype subsets becomes detailed to a level that is no longer meaningful for the global ancestry characterization of rice varieties at a population level.

To obtain an alternative indication for the optimal number of genomic clusters we evaluated our data set using Partitioning Around Medoids (PAM) with the *fviz_nbclust()* function from the ‘factoextra’ package (Kassambara and Mundt 2020) in R. We limited this analysis to *O. sativa* varieties. The PAM-based clustering tendency was assessed both with average silhouette scores (‘silhouette’) and gap statistic (‘gap_stat’) (Figures S13A+B). The former method showed a global optimum at K=3, and a local optimum at K=13. We used the *pam()* and *fviz_cluster()* functions to plot a multidimensional scaling plot (MDS) based on Euclidian distanced to visualize clusters for K=13, and *fviz_silhouette()* to generate silhouette plots for cluster validation (Figures S13C+D). The thirteen clusters discerned are (for accession-level details, see Data S5):

- focal population: ‘XI3 Cambodia’. XI3 varieties from Southeast Asia, East and West Africa
- focal population: ‘XI2 India’. All XI2 varieties
- focal population: ‘GJtrp Philippines’. GJtrp varieties from Southeast Asia, Madagascar and West Africa. In addition, the ‘GJtrp Maroon Alekisola’ and ‘GJtrp early-developed USA cultivars’ (e.g., *Rexoro*), and a subset of the ‘GJtrp Maroon Alekisola-admixed’ varieties
- focal population: ‘GJtrp Maroon core’. GJtrp ‘core’ varieties, as well as some other GJtrp varieties from Brazil, West Africa, Madagascar and the early-registered USA cultivars from Central and South America. In addition, all the ‘GJtrp Maroon long-awned’ varieties and a subset of the ‘GJtrp Maroon Alekisola-admixed’ varieties
- focal population: ‘Aus Bangladesh’. All circum-Aus varieties
- focal population: ‘XI3 Indonesia swamp’. All the ‘swamp’ ecotype varieties and a few other XI3 varieties from Southeast Asia. This group includes all the ‘XI3 Maroon swamp’ varieties
- focal population: ‘GJtrp Java’. All ‘GJtrp Java’ varieties
- focal population: ‘XI1A China’. All XI1A varieties
- focal population: ‘XI3 Laos hmonglike’. All XI3 ‘hmonglike’ varieties and a few XI3 varieties from Southeast Asia. This group includes the ‘XI3 Maroon Hmong’ variety
- focal population: ‘Bas India’. All Bas varieties
- focal population: ‘GJsbtrp Laos’. All GJsbtrp varieties
- focal population: ‘GJtmp South Korea’. All GJtmp varieties
- focal population: ‘GJtrp South Korea’. GJtrp varieties from Japan, South Korea and one from India

Taken together, we chose K=15 as our preferred local optimum for the ADMIXTURE model since it is the lowest *k*-value that includes the thirteen clusters also discerned with the PAM-based method (Table S4, Data S1D). The two additional ancestry components discerned at K=15 in ADMIXTURE maximize an ancestry component in the *O. glaberrima*, *O. barthii* and *O. longistaminata* varieties (these were excluded from the PAM-based analyses), and a second ecotype among Aus varieties, respectively. We grouped rice landraces that were collected in the same country and with a similar ADMIXTURE ancestry profile in a single population. As such, we followed a similar approach for population assignment as in human population genomic studies based on ancient DNA, which frequently rely on small population sizes or even single individuals as proxies for the ancestry of human groups associated with an archaeological context at a given area and moment in time (Sikora, et al. 2019; Mao, et al. 2021; Posth, et al. 2023; Villalba-Mouco, et al. 2023).

### Pairwise genomic distances

Genomic distances within *O. glaberrima*, *tropical japonica,* and *indica-3* rice subgroups were calculated between all pairs of rice landraces with formulation 1 − DST. DST was determined with the *--genome* and --*genome-full* options in PLINK v1.90b (Purcell, et al. 2007; Chang, et al. 2015), which is calculated: DST (IBS distance) = IBS2 + 0.5*IBS1) / (IBS0 + IBS1 + IBS2). Heat plots of 1-DST matrixes were made with the *pheatmap()* function from the pheatmap v1.0.12 package in R, which relies on the *hclust()* function from the stats package for hierarchical clustering.

### D- and F-statistics

For ancestry analyses on the population level, we used various *F_3_ and F_4_*-statistics methods (Reich, et al. 2009; Patterson, et al. 2012; Raghavan, et al. 2014) Standard errors (SE) were estimated using the 5cM block jackknife method (Busing, et al. 1999), as implemented in the *qpDstat* and *qp3Pop* programs in the ADMIXTOOLS v7.0 package (Patterson, et al. 2012). We define statistical significance as [-2.576 < Z < 2.576], corresponding to a 99% confidence interval (CI).

*Outgroup F_3_-statistics* (Raghavan, et al. 2014) of the form *f*_3_*(Outgroup; Pop_1_, Pop_2_)* measure the level of shared genomic drift between two groups of interest, *Pop_1_* and *Pop_2_*, relative to the *Outgroup* for SNPs ascertained in the *Outgroup*. The more closely related *Pop_1_* and *Pop_2_* are, the more positive the value for the *f_3_*-statistic becomes. We note that outgroup *f_3_*-statistics are mostly inconclusive in detecting what rice groups are the most closely related to the various GJtrp and XI3 Maroon rice groups. This is because of the large amount of ancestry that is shared among all *tropical japonica* or *indica(−3)* groups relative to the *Outgroup*. For example, in the outgroup *f_3_*-statistic *f_3_(O.barthii; GJtrp Maroon core, X)*, in which *X* are various worldwide GJtrp rice groups, any uniquely shared ancestry between ‘GJtrp Maroon core’ and *X* will be much smaller than the shared genetic drift between *O.barthii* and *tropical japonica* varieties in general. The relative increase in *f_3_*-value, due to the uniquely shared ancestry between ‘GJtrp Maroon core’ and *X*, is in general is too small to allow for comparison between various GJtrp groups *X*. As such, the 99% CIs typically overlap for these *f_3_*-outgroup statistics. To resolve what rice groups form the most closely related sister lineages to the Maroon rice groups, we instead use cladality *f_4_*-statistics.

*F_4_-statistics* (Reich, et al. 2009; Patterson, et al. 2012) are tests of treeness of four populations to estimate their correct phylogenetic relationship. *F_4_*-statistics of the form *f_4_(Outgroup, Pop_3_; Pop_1_, Pop_2_)* test the null-hypothesis that an AABB tree reflects the true tree topology of the four populations, and hence that *Pop_1_, Pop_2_* form a clade to the exclusion of *Pop_3_* and the *Outgroup.* If *Pop_1_* and *Pop_2_* form a true clade, both populations are expected to relate symmetrically to *Pop_3_* and the *Outgroup*. I.e., neither *Pop_1_* nor *Pop_2_* will show an excess of shared genetic drift to *Pop_3_* or the *Outgroup*. Hence, under the null-hypothesis for cladality (AABB tree) the *f_4_*-statistic will not significantly differ from *f_4_* = 0 (−2.576 < Z < 2.576, 99% CI). In contrast, if the *f_4_*-value differs significantly from zero this provides support for an alternative tree topology. The *f_4_*-statistic turns significantly negative (ABBA tree: Z < −2.576) when *Pop_1_* and *Pop_3_* share an excess of genetic drift, or when the *Outgroup* is not a true outgroup to the populations and shares an excess of drift with population *Pop_2._* Vice versa, the *f_4_*-statistic turns significantly positive (BABA tree, Z > 2.576) when *Pop_2_* and *Pop_3_* share an excess of genetic drift, or when the *Outgroup* is not a true outgroup to the populations and shares an excess of drift with population *Pop_1_*. In contrast to normalized D-statistics (Green, et al. 2010; Patterson, et al. 2012), the values of *f_4_*-statistics depend on the absolute allele frequencies of the SNPs used to calculate them (Lipson 2020). As such, *f_4_-*statistics are not only zero or nonzero but also carry quantitative information about amounts of shared drift between populations. Populations that share more drift yield longer intersecting (internal) branches and will have greater-magnitude *f_4_*-statistics associated with them. Another implication is that SNP sites that are invariable among the populations, i.e., ancestry that these populations have in common, will shrink the *f_4_*-statistics towards zero (Lipson 2020). The same *f_4_-*statistic for a rooted tree can also be used as an admixture *f_4_*-statistic to test for admixture of *Pop_3_* into *Pop_1_* or *Pop_2_* after the divergence between *Pop_1_* and *Pop_2_*, and under the assumption that *Pop_1_* is (a close proxy to) the ancestry progenitor of *Pop_2_* or vice versa.

*Admixture f_3_-statistics* (Reich, et al. 2009) of the form *f*_3_(*Target; Pop_1_, Pop_2_)* test whether the allele frequencies of the *Target* population are intermediate between that of populations *Pop_1_* and *Pop_2_*, and hence if the *Target* population can be considered as admixed between *Pop_1_* and *Pop_2_*. Allele frequencies of the *Target* population that are intermediate between *Pop_1_* and *Pop_2_* result in drift on the path on internal branches of the population tree and force the *f_3_*-statistic negative. Post-admixture genetic drift in the *Target* population tends to make the *f*_3_-statistic less negative (Peter 2016), making this a more conservative test for admixture than an *f*_4_-statistic.

## Supplementary Material

Supplementary data are available at Molecular Biology and Evolution online.

## Acknowledgments

We would like to express our gratitude to the Maroon farmers in Suriname and French Guiana who shared their knowledge with us and provided rice varieties. In addition, we thank Paul Richards and Paul Struik for sharing their anthropological and historical perspectives on the genomic results, and Siva Selvanayagam for providing advice on genotype QC.

## Author contributions

Conceptualization: TvA, ES. Funding acquisition: TvA, ES, RvV. Sample collection and ethnobotanical surveys: NP, TvA. Sample propagation, phenotyping and storage in germplasm bank: JTA. Genomic lab work: FB, MvdL. Genomic analyses: MvdL. Data integration: MvdL, TvA, NP, HM, ES, RvV and JTA. Data visualization: MvdL. Supervision: TvA, ES, HM. Writing – original draft: MvdL, TvA, ES. Writing – review and editing: all authors

## Data Availability

Genomic data (BAM format) are available through the Sequence Read Archive (accession number PRJNA1019116). The genotype data for the 110k SNPs can be downloaded via XXX.

## References

1. Agrama HA, Yan W, Jia M, Fjellstrom R, McClung AM. 2010. Genetic structure associated with diversity and geographic distribution in the USDA rice world collection. Natural Science. 2(4):247–291. doi: 10.4236/ns.2010.24036

2. Alexander DH, Novembre J, Lange K. 2009. Fast model-based estimation of ancestry in unrelated individuals. Genome Research. 19(9):1655–1664. doi: 10.1101/gr.094052.109

3. Busing FMTA, Meijer E, Leeden RVD. 1999. Delete-m Jackknife for Unequal m. Statistics and Computing. 9:3–8. doi: 10.1023/A:1008800423698

4. Carney JA. 1998. The Role of African Rice and Slaves in the History of Rice Cultivation in the Americas. Human Ecology. 26(4):525–545. doi: 10.1023/A:1018716524160

5. Carney JA. 2002. Black Rice. Cambridge and London: Harvard University Press.

6. Carney JA. 2005. Rice and memory in the age of enslavement: Atlantic passages to Suriname. Slavery & Abolition. 26(3):325–348. doi: 10.1080/01440390500319562

7. Chang CC, Chow CC, Tellier LC, Vattikuti S, Purcell SM, Lee JJ. 2015. Second-generation PLINK: rising to the challenge of larger and richer datasets. GigaScience. 4:7. doi:10.1186/s13742-015-0047-8

8. Chen S, Zhou Y, Chen Y, Gu J. 2018. fastp: an ultra-fast all-in-one FASTQ preprocessor. Bioinformatics. 34(17):i884–i890. doi: 10.1093/bioinformatics/bty560

9. Clarkin PF. 2005. Hmong resettlement in French Guiana. Hmong Studies Journal. 6:1–27.

10. Czech L, Exposito-Alonso M. 2022. grenepipe: a flexible, scalable and reproducible pipeline to automate variant calling from sequence reads. Bioinformatics. 38(20):4809–4811. doi: 10.1093/bioinformatics/btac600

11. Dragtenstein F, 2004. “De ondraaglijke stoutheid der wegloopers”: marronage en koloniaal beleid in Suriname, 1667-1768. Bijdragen en mededelingen betreffende de geschiedenis der Nederlanden. 119(1):99. doi: 10.18352/bmgn-lchr.5985

12. Dragtenstein F. 2017. Van Elmina naar Paramaribo: de slavenhaler. Zutphen: Walburg Press.

13. Eltis D, Richardson D. 1995. Atlas of the Transatlantic Slave Trade. New Haven: Yale University Press

14. Fleury M. 1994. Impact de la traite des esclaves sur la phytogéographie: exemple chez les Aluku (Boni) de Guyane française. Journal d’Agriculture Traditionnelle et de Botanique Appliquée 36:113–137. doi: 10.3406/jatba.1994.3537

15. Gaio D, Anantanawat K, To J, Liu M, Monahan L, Darling AE. 2022. Hackflex: low-cost, high-throughput, Illumina Nextera Flex library construction. Microbial Genomics. 8(1):000744. doi: 10.1099/mgen.0.000744

16. Garrison EP, Marth GT. 2012. Haplotype-based variant detection from short-read sequencing. arXiv. 1207.3907. doi: 10.48550/arXiv.1207.3907

17. Gilbert E. 2015. Asian Rice in Africa: Plant Genetics and Crop History. In: Schäfer D, Fields-Black EL, Bray F, Coclanis PA, editors. Rice: Global Networks and New Histories. Cambridge: Cambridge University Press. p. 212-228

18. Green RE, Krause J, Briggs AW, Maricic T, Stenzel U, Kircher M, Patterson N, Li H, Zhai W, Fritz MH, et al. 2010. A draft sequence of the Neandertal genome. Science. 328(5979):710–722.

19. Gutaker RM, Groen SC, Bellis ES, Choi JY, Pires IS, Bocinsky RK, Slayton ER, Wilkins O, Castillo CC, Negrão S, et al. 2020. Genomic history and ecology of the geographic spread of rice. Nature Plants. 6:492–502. doi: 10.1038/s41477-020-0659-6

20. Han B, Xue Y. 2003. Genome-wide intraspecific DNA-sequence variations in rice. Current Opinion in Plant Biology. 6(2):134–138. doi: 10.1016/s1369-5266(03)00004-9

21. Hoefte R. 1998. In Place of Slavery: A Social History of British Indian and Javanese Laborers in Suriname. Gainesville: University Press of Florida

22. Hoogbergen WSM, Polimé T. 2002. Oostelijk Suriname 1986-2002. In: OSO. Tijdschrift voor Surinaamse taalkunde, letterkunde en geschiedenis. Jaarkgang 21. Nijmegen: Stichting Instituut ter Bevordering van de Surinamistiek. p. 225–242

23. Huang X, Kurata N, Wei X, Wang Z-X, Wang A, Zhao Q, Zhao Y, Liu K, Lu H, Li W, et al. 2012. A map of rice genome variation reveals the origin of cultivated rice. Nature. 490:497–501. doi: 10.1038/nature11532

24. Iskandar J, Ellen R. 1999. In situ conservation of rice landraces among the Baduy of west Java. Journal of Ethnobiology. 19(1):97–125.

25. Kassambara A, Mundt F. 2020. Factoextra: Extract and Visualize the Results of Multivariate Data Analyses. R Package Version 1.0.7. https://CRAN.R-project.org/package=factoextra

26. Kawahara Y, de la Bastide M, Hamilton JP, Kanamori H, McCombie WR, Ouyang S, Schwartz DC, Tanaka T, Wu J, Zhou S, et al. 2013. Improvement of the Oryza sativa Nipponbare reference genome using next generation sequence and optical map data. Rice. 6(1):4. doi: 10.1186/1939-8433-6-4

27. Larranaga N, van Zonneveld M, Hormaza JI. 2021. Holocene land and sea-trade routes explain complex patterns of pre-Columbian crop dispersion. New Phytologist. 229:1768–1781. doi: 10.1111/nph.16936

28. Li H, Handsaker B, Wysoker A, Fennell T, Ruan J, Homer N, Marth G, Abecasis G, Durbin R, 1000 Genome Project Data Processing Subgroup. 2009. The Sequence Alignment/Map format and SAMtools. Bioinformatics. 25(16):2078–2079. doi: 10.1093/bioinformatics/btp352

29. Lipson M. 2020. Applying f4-statistics and admixture graphs: Theory and examples. Molecular Ecology Resources. 20(6):1658–1667. doi: 10.1111/1755-0998.13230

30. Lu H, Redus M, Coburn J, Rutger J, McCouch S, Tai T. 2005. Population Structure and Breeding Patterns of 145 U.S. Rice Cultivars Based on SSR Marker Analysis. Crop Science. 45: 66–76. doi: 10.2135/cropsci2005.0066

31. Maat H. 2015. Commodities and Anti-Commodities: Rice on Sumatra 1915–1925. In: Schäfer D, Fields-Black EL, Bray F, Coclanis PA, editors. Rice: Global Networks and New Histories. Cambridge: Cambridge University Press. p. 335–354.

32. Maat H, van Andel TR. 2018. The history of the rice gene pool in Suriname: Circulations of rice and people from the eighteenth century until late twentieth century. Historia Agraria. 75:69–91. doi: 10.26882/histagrar.075e04m

33. Mao X, Zhang H, Qiao S, Liu Y, Chang F, Xie P, Zhang M, Wang T, Li M, Cao P, et al. 2021. The deep population history of northern East Asia from the Late Pleistocene to the Holocene. Cell. 184(12):3256–3266.e13. doi: 10.1016/j.cell.2021.04.040

34. McKenna A, Hanna M, Banks E, Sivachenko A, Cibulskis K, Kernytsky A, Garimella K, Altshuler D, Gabriel S, Daly M, DePristo MA. 2010. The Genome Analysis Toolkit: a MapReduce framework for analyzing next-generation DNA sequencing data. Genome Research. 20(9):1297–1303. doi: 10.1101/gr.107524.110

35. Meyer RS, Choi JY, Sanches M, Plessis A, Flowers JM, Amas J, Dorph K, Barretto A, Gross B, Fuller DQ, et al. 2016. Domestication history and geographical adaptation inferred from a SNP map of African rice. Nature Genetics. 48(9):1083–1088. doi: 10.1038/ng.3633

36. Mokuwa A, Nuijten E, Okry F, Teeken B, Maat H, Richards P, Struik PC. 2013. Robustness and Strategies of Adaptation among Farmer Varieties of African Rice (Oryza glaberrima) and Asian Rice (Oryza sativa) across West Africa. PLoS One. 8(3):e34801. doi: 10.1371/journal.pone.0034801

37. Nawani S. 2013. The Portuguese in archipelago Southeast Asia (1511-1666). Proceedings of the Indian History Congress. 74:703–708

38. Patterson N, Price AL, Reich D. 2006. Population structure and eigenanalysis. PLoS Genetics. 2(12):e190. doi: 10.1371/journal.pgen.0020190

39. Patterson N, Moorjani P, Luo Y, Mallick S, Rohland N, Zhan Y, Genschoreck T, Webster T, Reich D. 2012. Ancient Admixture in Human History. Genetics. 192(3):1065–1093. doi: 10.1534/genetics.112.145037

40. Peter BM. 2016. Admixture, Population Structure, and F-Statistics. Genetics. 202(4):1485–1501. doi: 10.1534/genetics.115.183913

41. Pinas N, van de Loosdrecht M, Maat H, van Andel T. 2023. Vernacular Names of Traditional Rice Varieties Reveal the Unique History of Maroons in Suriname and French Guiana. Economic Botany. 77:117–134. doi: 10.1007/s12231-023-09571-0

42. Posth C, Yu H, Ghalichi A, Rougier H, Crevecoeur I, Huang Y, Ringbauer H, Rohrlach AB, Nägele K, Villalba-Mouco V, et al. 2023. Palaeogenomics of Upper Palaeolithic to Neolithic European hunter-gatherers. Nature. 615:117–126. doi: 10.1038/s41586-023-05726-0

43. Price R. 1991. Subsistence on the plantation periphery: Crops, cooking, and labour among eighteenth-century Suriname maroons. Slavery & Abolition. 12(1):107–127. doi: 10.1080/01440399108575025

44. Price S. 1993. Co-wives and Calabashes: Michigan: University of Michigan Press. doi: 10.3998/mpub.7914

45. Price R. 2018. Maroons in Guyane: Getting the Numbers Right. New West Indian Guide. 92:275–283. doi: 10.1163/22134360-09203001

46. Purcell S, Neale B, Todd-Brown K, Thomas L, Ferreira MAR, Bender D, Maller J, Sklar P, de Bakker PIW, Daly MJ, Sham PC. 2007. PLINK: A Tool Set for Whole-Genome Association and Population-Based Linkage Analyses. The American Journal of Human Genetics. 81(3):559–575. doi: 10.1086/519795.

47. Quinlan AR, Hall IM. 2010. BEDTools: a flexible suite of utilities for comparing genomic features. Bioinformatics. 26(6):841–842. doi: 10.1093/bioinformatics/btq033

48. Raghavan M, Skoglund P, Graf KE, Metspalu M, Albrechtsen A, Moltke I, Rasmussen S, Stafford Jr TW, Orlando L, Metspalu E, et al. 2014. Upper Palaeolithic Siberian genome reveals dual ancestry of Native Americans. Nature. 505(7481):87–91. doi: 10.1038/nature12736

49. Reich D, Thangaraj K, Patterson N, Price AL, Singh L. 2009. Reconstructing Indian population history. Nature. 461(7263):489–494. doi: 10.1038/nature08365

50. Rutger JN, Mackill DJ. 2008. Application of Mendelian genetics in rice breeding. In: Rice Genetics IV. p.27–38. doi: 10.1142/9789812814296_0002

51. Scott MF, Botigué LR, Brace S, Stevens CJ, Mullin VE, Stevenson A, Thomas MG, Fuller DQ, Mott R. 2019. A 3,000-year-old Egyptian emmer wheat genome reveals dispersal and domestication history. Nature Plants. 5(11):1120–1128. doi: 10.1038/s41477-019-0534-5

52. Sikora M, Pitulko VV, Sousa VC, Allentoft ME, Vinner L, Rasmussen S, Margaryan A, de Barros Damgaard P, de la Fuente C, Renaud G, et al. 2019. The population history of northeastern Siberia since the Pleistocene. Nature. 570(7760):182–188. doi: 10.1038/s41586-019-1279-z

53. Stahel G. 1944. De nuttige planten van Suriname. In: Bulletin Landbouwproefstation in Suriname; no. 59. Paramaribo: Departement Landbouwproefstation in Suriname. The 3,000 rice genomes project. 2014. The 3,000 rice genomes project. GigaScience 3:7. doi: 10.1186/2047-217X-3-7

54. Thoden van Velzen HUE. 2022. Prophets of doom: a history of the Okanisi Maroons in Suriname. Leiden: Brill.

55. van Andel TR. 2010. African Rice (Oryza glaberrima Steud.): Lost Crop of the Enslaved Africans Discovered in Suriname. Economic Botany. 64(1):1–10. doi: 10.1007/s12231-010-9111-6

56. van Andel TR, Meyer RS, Aflitos SA, Carney JA, Veltman MA, Copetti D, Flowers JM, Havinga RM, Maat H, Purugganan MD, et al. 2016. Tracing ancestor rice of Suriname Maroons back to its African origin. Nature Plants. 2:16149. doi: 10.1038/nplants.2016.149

57. van Andel TR, van der Velden A, Reijers M. 2016. The ‘Botanical Gardens of the Dispossessed’ revisited: richness and significance of Old World crops grown by Suriname Maroons. Genetic Resources and Crop Evolution. 63:695–710. doi: 10.1007/s10722-015-0277-8

58. Van Andel TR, Veltman MA, Bertin A, Maat H, Polime T, Hille Ris Lambers D, Tjoe Awie J, De Boer H, Manzanilla V. 2019. Hidden Rice Diversity in the Guianas. Frontiers in Plant Science. 10:1161. doi: 10.3389/fpls.2019.01161

59. van Andel TR, Maat H, Pinas N. 2023. Maroon Women in Suriname and French Guiana: Rice, Slavery, Memory. Slavery & Abolition. 1–25. doi: 10.1080/0144039X.2023.2228771

60. Vasimuddin M, Misra S, Li H, Aluru S editors. 2019 IEEE International Parallel and Distributed Processing Symposium (IPDPS). 20-24 May 2019

61. Vaughn JN, Korani W, Stein JC, Edwards JD, Peterson DG, Simpson SA, Youngblood RC, Grimwood J, Chougule K, Ware DH, et al. 2021. Gene disruption by structural mutations drives selection in US rice breeding over the last century. PLoS Genetics. 17(3):e1009389. doi: 10.1371/journal.pgen.1009389

62. Veltman MA, Flowers JM, van Andel TR, Schranz ME. 2019. Origins and geographic diversification of African rice (Oryza glaberrima). PLoS One. 14(3):e0203508. doi: 10.1371/journal.pone.0203508

63. Villalba-Mouco V, van de Loosdrecht MS, Rohrlach AB, Fewlass H, Talamo S, Yu H, Aron F, Lalueza-Fox C, Cabello L, Cantalejo Duarte P, et al. 2023. A 23,000-year-old southern Iberian individual links human groups that lived in Western Europe before and after the Last Glacial Maximum. Nature Ecology & Evolution. 7(4):597–609. doi: 10.1038/s41559-023-01987-0

64. Wang W, Mauleon R, Hu Z, Chebotarov D, Tai S, Wu Z, Li M, Zheng T, Fuentes RR, Zhang F, et al. 2018. Genomic variation in 3,010 diverse accessions of Asian cultivated rice. Nature. 557(7703):43–49. doi: 10.1038/s41586-018-0063-9

65. Wood PH. 1975. Black majority: Negroes in colonial South Carolina from 1670 through the Stono Rebellion. New York: WW Norton.

66. Zerega NJ, Ragone D, Motley TJ. 2004. Complex origins of breadfruit (Artocarpus altilis, Moraceae): implications for human migrations in Oceania. American Journal of Botany. 91(5):760–766. doi: 10.3732/ajb.91.5.760

67. Zeven AC. 1998. Landraces: A review of definitions and classifications. Euphytica. 104:127–139. doi: 10.1023/A:1018683119237

